# Polypide anatomy of hornerid bryozoans (Stenolaemata: Cyclostomatida)

**DOI:** 10.1101/2021.03.09.433978

**Authors:** Yuta Tamberg, Peter B. Batson, Ruth Napper

## Abstract

Bryozoans are small colonial coelomates whose colonies are made of individual modules (zooids). Like most coelomate animals, bryozoans have a characteristic body wall composition, including epidermis, extracellular matrix (ECM) and coelothelium, all pressed together. The order Cyclostomatida, however, presents the most striking deviation, in which the ECM and the corresponding coelothelium underlying major parts of the skeletal wall epidermis are “;peeled off” to form an independent membranous sac. The polypide anatomy and ultrastructure of this group is best known from one family, the Crisiidae (Articulata). Here we examined four species from the phylogenetically and ecologically contrasting family Horneridae (Cancellata) from New Zealand. Here we provide the first detailed ultrastructural examination of the hornerid polypide, including tentacles, mouth region, digestive system and the funiculus. We were able to trace continuity and transitions of cell and ECM layers throughout the whole polypide. In addition we identified that the funiculus is a lumen-free ECM cord with two associated muscles, disconnected from interzooidal pores. While agreeing with the general cyclostomate body plan, hornerids have some unique traits that make them worthy of additional study.

**Highlights:** Hornerids share a general cyclostomate body plan. The frontal tentacle ECM transitions into oral sphincter ECM, the abfrontal lophophore ECM becomes a septum between coelomic compartments, and the funuculus is a solid ECM cord supplied with muscles.

## 1. Introduction

Bryozoans are small colonial benthic invertebrates living in marine and freshwater environments. The phylum contains three classes (Phylactolaemata, Gymnolaemata, Stenolaemata). Bryozoan colonies are made of individual units, or modules, called zooids, linked with colony-wide elements. The feeding zooids comprise an outer skeletal wall (ectocyst), the living layer or layers of tissues attached to and underlying the skeleton (endocyst), and the polypide—the retractable soft-body part of the zooid which includes all its organs.

Bryozoans are coelomic animals with two or three compartments in their body cavities, partially confluent or completely separated (Shunatova & Tamberg, 2019). In Phylactolaemata these compartments include the coeloms of the epistome, lophophore, and trunk, in Gymnolaemata—the lophophore and trunk coeloms, connected by narrow ciliated ducts. In Stenolaemata there is a lophophore coelom and a modified trunk cavity with two portions. Stenolaemate polypides, unlike other bryozoans, are surrounded by a membranous sac, made of a single coelothelial layer and the ECM, and supplied with delicate muscles (Nielsen & Pedersen, 1979), whereas the epidermal layer is resting against the skeletal wall. Thus, their main body cavity is split into two parts: the exosaccal cavity outside the membranous sac, and the endosaccal cavity (true coelom) inside it (Figure 1).

**Figure 1.**
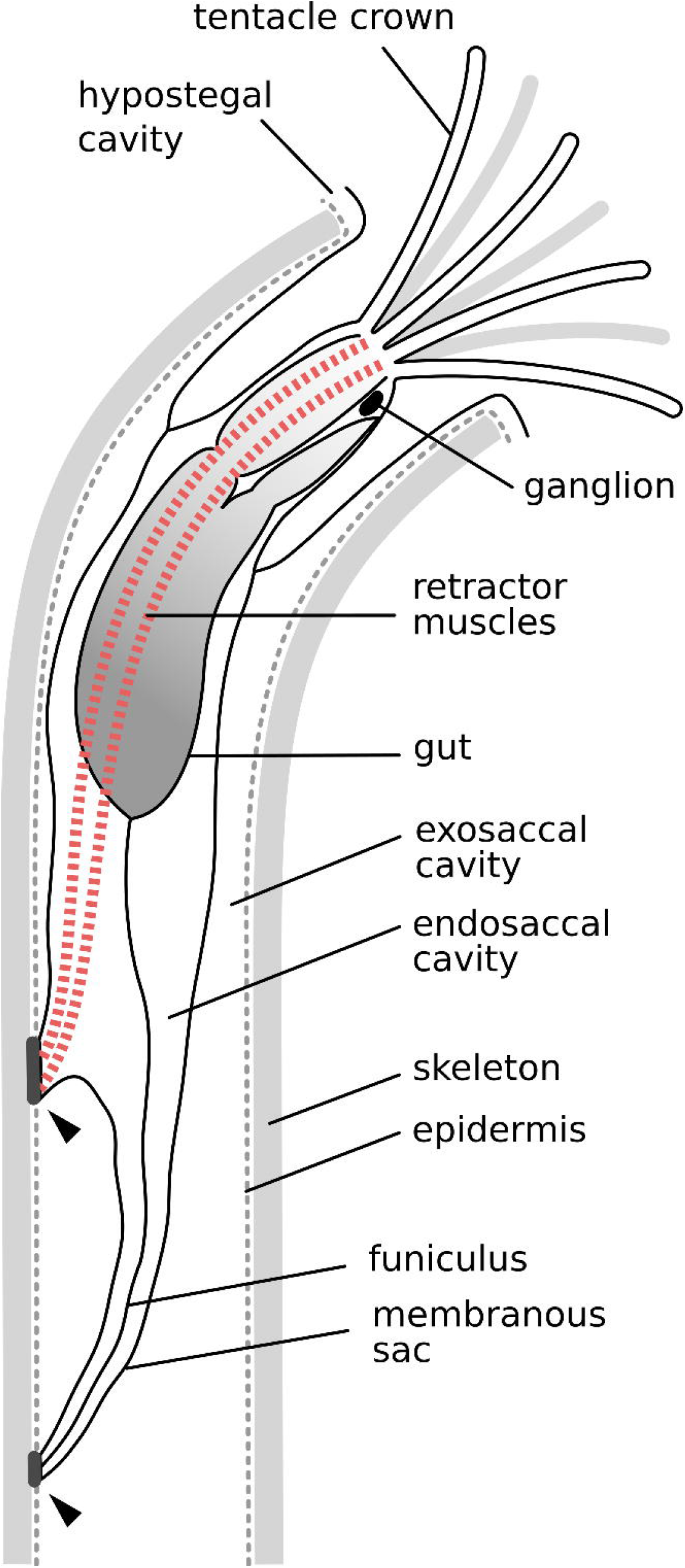
Schematic drawing of the protruded hornerid zooid. Arrowheads, membranous sac-to-cystid attachments shared by all studied species

The coelomic lining of the tentacles, the lophophore and the tentacle sheath is similar in all bryozoan classes, whereas the trunk lining differs across these groups. Phylactolaemata have an unbroken coelothelium, cheilostomates within Gymnolaemata have lost parts of the ECM and peritoneum, and Stenolaemata have ECM and peritoneum detached from the trunk epidermis in the form of the membranous sac.

In stenolaemates the membranous sac is not completely detached from the cystid. In a wide range of studied species, it has been shown to be anchored to the skeleton proximally at the origin points of the polypide retractor muscles and the funiculus; in addition, there may be various polypide-cystid attachments in the distal part of the zooid (e.g. Borg, 1926; Boardman and McKinney, 1985; Boardman 1998; Nielsen, 1970; Nielsen and Pedersen, 1979; Schäfer, 1985; Shunatova and Tamberg, 2019; Schwaha et al., 2018). All such structures provide stability to the polypide during protrusion and retraction.

The Cyclostomatida is the only living order in the exclusively marine class Stenolaemata. Our understanding of the anatomy of cyclostomate polypides is mostly based on light-microscopic studies by Borg (1926), Boardman (1983, 1998), Boardman et al. (1992), Schäfer (1985), Nielsen (1970) and recently Schwaha et al. (2018). Only a few publications have addressed cyclostomate ultrastructure using electron microscopy: Nielsen and Pedersen (1979), Carle and Ruppert (1983), Nielsen and Riisgard (1998), Shunatova & Nielsen (2002), Nielsen (2013), Temereva and Kosevich (2018) and Shunatova and Tamberg (2019). In addition, some confocal laser scanning microscopic studies have examined nervous and muscular systems of these animals (Schwaha et al., 2018, Temereva & Kosevich, 2018, Worsaae et al., 2018).

Although all these studies provided major insights into cyclostomate anatomy, they cover a relatively narrow taxonomic range. A single genus, *Crisia* (suborder Articulata), is the most-studied cyclostomate; six out of nine recent electron microscopic and confocal studies listed above explore *Crisia eburnea* or *Crisia elongata*. Variations of Borg’s (1926) and Nielsen and Pedersen’s (1979) *Crisia* autozooid schematics remain the standard depiction of generalized cyclostomate anatomy. Representatives of Tubuliporidae, Lichenoporidae and Cinctiporidae are much less studied, and other families remain virtually unexplored at the ultrastructural level.

The cancellate cyclostomates in the family Horneridae differ in several key respects from *Crisia*. The latter is a delicate, ephemeral, subtidal cyclostomate, whereas hornerids are heavily calcified, long-lived, and typically occur in oceanic environments at shelf depths and deeper. Recent molecular phylogeny has confirmed that the Cyclostomatida is monophyletic and that, within this order, hornerids are widely separated from articulates (Waeschenbach et al., 2012). For these reasons, a comparison of hornerids with *Crisia* can highlight the range of anatomical variation across the Cyclostomatida.

In this paper we explore polypide anatomy and ultrastructure of four hornerid species from southern New Zealand using light and electron microscopic methods. In particular, we trace various epithelial layers of the polypide at the ultrastructural level and compare them to other bryozoans, in order to develop a deeper, more multidimensional understanding of the anatomy of this group. This understanding could lead to improved reconstructions of soft body parts of extinct stenolaemates (see Tamberg & Smith, 2020).

## 2. Material and Methods

We examined four hornerid species from two genera: *Hornera robusta* MacGillivray, 1883, *Hornera* sp. 1, *Hornera* sp. 2 and Horneridae gen. sp. 3. The first three species, *H. robusta, H*. sp. 1 and *H*. sp. 2, were collected annually from 2016 to 2019 from mid-continental shelf off Otago, New Zealand (90 m depth, 45° 47.89’ S, 170° 54.5’ E; see Batson and Probert 2000). *Hornera* sp. 2 was also sampled in 2018 near Stewart Island (58 m and 77 m; 46°54.87’ S, 168°13.06’ E and 47°07.70’ S, 168°10.79’ E respectively). The last species, Horderidae gen. sp. 3 (planar and curved forms) was obtained in 2016 near Snares Islands (151 m; 47° 43.20’ S, 167° 1.44’ E). Upon collection some colonies were fixed immediately, while others were transported alive to the lab.

Living colonies of *H. robusta* and *H*. sp. 2 were kept in flow tanks in an isothermic room at ∼13°C, where they were left to recover from dredging for 3–8 days. Throughout this time the animals were constantly supplied with a mixture of natural particles and cultured algae *Rhodomonas salina* and *Dunaliella tertiolecta* using a drip feeder system. Colonies were than examined under the microscope, and fully recovered feeding ones were relaxed with an isotonic solution of magnesium chloride (∼7.5%) mixed 1:1 with sea water and fixed. Colonies of *H*. sp. 1 and Horneridae gen. sp. 3 were fixed immediately upon collection, so that all polypides are in a retracted state.

We treated the fixed colonies following three processing pathways: paraffin-embedded material was used for histological examination, epoxy-resin embedded tissue was used for semi-thin sections and TEM, and some material was also specifically processed and embedded in resin for imaging with serial block-face SEM (SBF SEM).

For paraffin sections, we fixed the colonies of *H. robusta* with 4% formalin in sea water, washed, dehydrated through the graded ethanol series and embedded in paraffin (Peterfi method). The 5 µm thick sections were stained with hematoxylin-eosin.

To obtain semi-thin and ultrathin sections, we fixed samples with 2.5% glutaraldehyde solution in 0.1 M PBS supplemented with sucrose to reach 990 mOsmol, and processed using standard TEM protocols. Material was rinsed in 0.1 M PBS with sucrose (990 mOsmol), decalcified with EDTA and transferred into 1% OsO_4_ for 1.5 hours. After osmication, the material was washed, dehydrated through a graded ethanol series and pure acetone, and embedded in Embed 812 epoxy resin. Sectioning was done with a diamond knife on Leica EM UC7 ultramicrotome. Resulting series of semi-thin sections (1 µm thick) were stained with toluidine blue and imaged with a light microscope. Ultrathin sections (80–90 nm) were stained with uranyl acetate and lead citrate and imaged with JEOL JEM 1400 and JEOL 2200FS electron microscopes.

Two colonies of *H. robusta* were fixed with glutaraldehyde and processed for SBF SEM (protocol modified from Deerinck et al., 2010). The samples were washed with buffer, pre-treated with 0.1% tannic acid in buffer for 45 minutes, washed with the same buffer, and post-fixed with 1.5% potassium ferrocyanide and 1% osmium tetroxide mixture on ice for 1.5 hours. All following washes were done with distilled water. The samples were placed in 1% thiocarbohydrazide solution for 30 minutes at 60°C, and in 2% OsO_4_ for 60 minutes. Samples were then stained consecutively with 1% uranyl acetate overnight and with lead aspartate solution for 30 minutes at 60°C. Three 5–7 minute washes were used between each reagent. After the final wash samples were dehydrated through graded ethanol series and embedded in Durcupan, ACM resin (Electron Microscopy Sciences, USA). SBF SEM sectioning and imaging was done with ZEISS Sigma VP 3View (Sydney University) and Thermo Fisher Apreo with VolumeScope Serial Block Face system (Queensland University) electron microscopes.

Electron microscope images are partially represented as negatives converted by ImageMagick 6.9. (The ImageMagick Development Team 2021), but not altered otherwise. Tiled montages were aligned with Etomo element of IMOD software (Kremer et al., 1996). All measurements were done from microphotographs using Inkscape 0.92 (Inkscape project 2017).

## 3. Results

All hornerid species have tubular zooidal chambers up to 2 mm long, 60–100 µm in internal diameter, narrowing sharply at the proximal end. Regardless of the size and age of the zooid, the polypide invariably occupies the distal-most position, moving 300–500 µm up and down the zooid tube when protruding and retracting. This indicates that both examined genera undergo progressive polypide cycles (*sensu* Boardman, 1998). There are at least two points where the polypide is attached to the zooid wall: the origin of the retractor muscles (∼450 µm from the aperture) and the funiculus (additional ∼350 μm). In both Horneridae gen. sp. 3 forms the polypide is additionally anchored to the skeletal wall by distal ligaments.

When the polypide is protruded, nine 200–300 µm long tentacles curve outwards in a trumpet-shaped crown. The tentacles protrude from the aperture for 90%–100% of their length. When retracted, tentacles are straightened and arranged in a cylinder.

The general anatomy of the studied species is relatively uniform and broadly resembles that described in other Cyclostomatida (Figure 1). In the morphological descriptions, given below, we mostly provide new information or address the points of difference.

### 3.1 Tentacles

Tentacles are roughly triangular in cross-section, frontal–abfrontal distance is ∼17 µm; maximum lateral width is ∼12 µm (including cuticle). Organization of the tentacles follows the standard bryozoan plan: a tube of ECM is surrounded by outer epithelial cells arranged in regular vertical rows, and lined internally with coelothelium. The coelomic cavity is seen as a series of lacunae in both protruded and retracted tentacles.

The outer epithelial layer includes: [1] an unpaired midline frontal A-cell, [2] two lateral– frontal B-cells on either side, [3] two frontal–lateral C-cells, [4] two lateral D-cells, [5] two abfrontal–lateral E-cells and (6) an intermittent row of abfrontal F-cells (Figure 2a,b).

**Figure 2.**
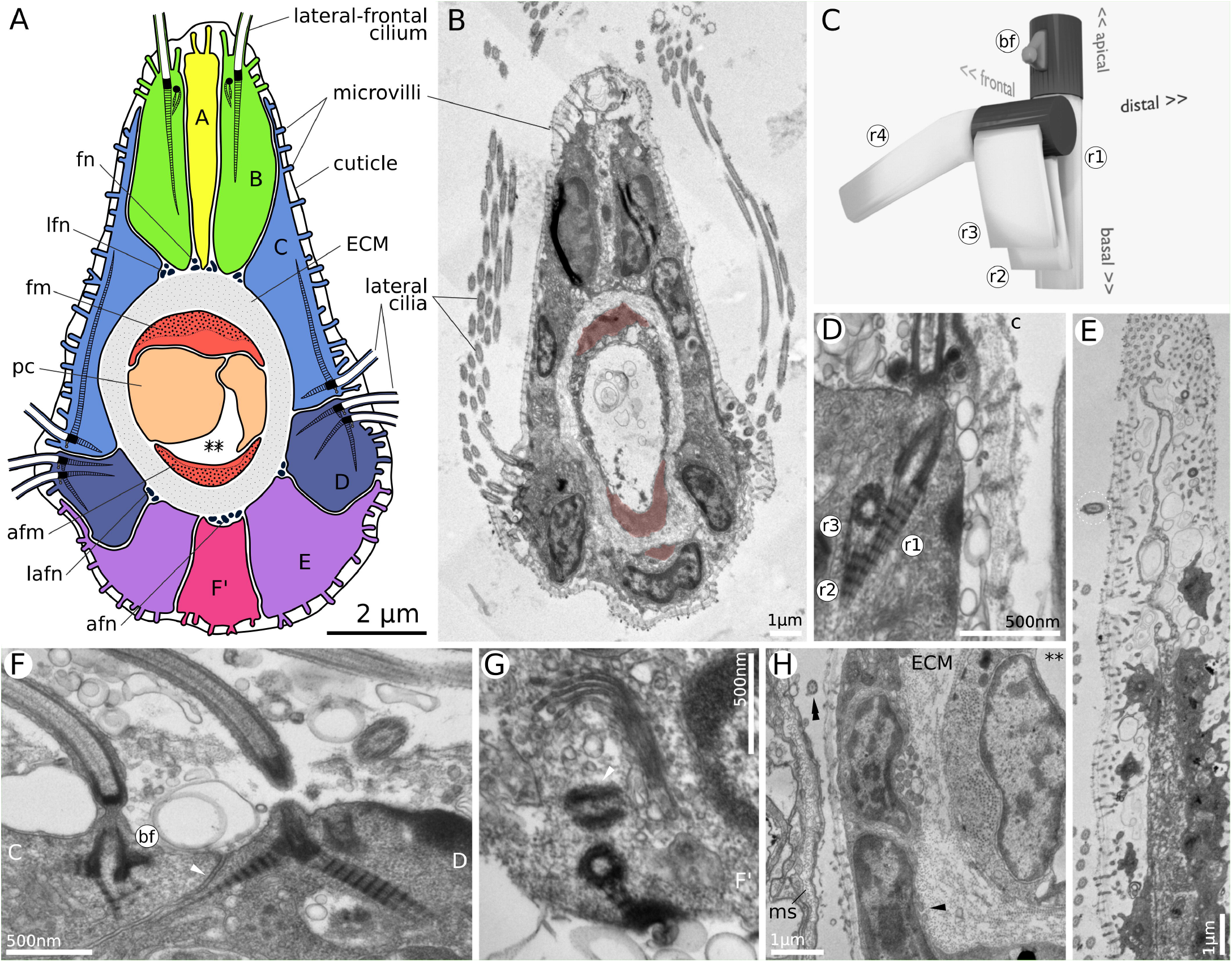
Ultrastructure of hornerid tentacles. (a) Schematic drawing of a tentacle cross-section. (b) Cross-section through the middle part of a protruded tentacle (*H*. sp. 2). (c) 3D model of the lateral-frontal ciliary basal complex. (d) Transverse section of a lateral-frontal cell showing ciliary basal apparatus (*H. robusta*, tentacle midline on the left). (e) Oblique frontal section of the tentacle showing frontal microvilli and a double row of lateral-frontal cilia (*H*. sp. 2). (f) Transverse section of lateral cells showing the basal apparatus of basal cilia and cell contacts (*H. robusta*, frontal side of the tentacle on the left). (g) Cross-section of an abfrontal cell showing basal apparatus of the abfrontal cilia (*H. robusta*, apical surface at the bottom). (h) Cross-section showing the abfrontal part of the tentacle, abfrontal and abfrontal-lateral nerves, and abfrontal cilium (*H. robusta*, abfrontal surface on the left). A, frontal cell of tentacle epidermis; B, lateral-frontal cell; C, frontal-lateral cell; D, lateral cell; E, abfrontal-lateral cell; F’, intermittent abfrontal cell; afm, abfrontal tentacle muscle; afn, abfrontal nerve; c, cuticle; ECM, extracellular matrix; fm, frontal tentacle muscle; fn, frontal nerve; lafn, abfrontal-lateral nerve; lfn, lateral-frontal nerve; pc, peritoneal cell; **, lophophore coelom; black arrowhead, abfrontal-lateral nerve; double back arrowhead, abfrontal cilium; white arrowhead, septate junction

A-cells are oval in cross-section, elongated (∼8 μm) along the main tentacle axis, usually with electron-light cytoplasm. In full agreement with previous observations, A-cells never bear cilia. In *H. robusta* the cytoplasm of A-cells often contains numerous vesicles near the apical tip. In *Hornera* species 1 and 2 the tips of A-cells bear long (1.2–2.2 μm) branching microvilli, while in hornerid gen. sp. 3 they are much shorter (∼0.5 μm).

B-cells are of similar size and shape in cross-section, but are much shorter (∼2 μm) in proximal–distal direction. These are monociliated cells with deeply lobed nuclei and complicated basal complexes including two kinetosomes and four cross-striated rootlets (r1-4 on Figure 2c,d). On each tentacle, lateral-frontal cilia form two closely-spaced rows (Figure 2e). The basal complexes in B-cells on the right and left of the tentacle are mirror-images of each other. The main kinetosome is attached to the apical cell membrane by several short microtubules. In addition, it has one axial cross-striated rootlet (r1) and a basal foot. Both the basal foot and the additional kinetosome are located on the frontal side, i.e., closer to the midline of the tentacle. An additional kinetosome gives off three rootlets: two axial ones (r2, 3) and a diagonal one (r4). All axial rootlets (r1, 2, and 3) run close to each other, straight towards the base of the cell, whereas the diagonal rootlet extends in a proximal– basal direction.

C-cells cover most of the frontal–lateral surfaces of the tentacles with their elongated flaps pointing forward. They come up nearly to the apical tips of the B-cells but form near-vertical abfrontal borders with the D-cells. In proximal-distal direction C-cells extend for ∼5 μm. C-cells bear numerous long cilia (∼15 µm) which form the frontal half of the lateral ciliary row (Figure 2b). Each cilium has a single basal body with a basal foot and two cross-striated rootlets (Figure 2f, cell on the left). The axial rootlet extends basally and abfrontally at a steep angle, whereas a very long lateral rootlet runs underneath the cell membrane in the frontal direction into the cytoplasmic flap. As in other bryozoans, the basal foot always points abfrontally, in the direction of the active ciliary stroke. The microvilli of C-cells are usually short (∼0.2 μm), except for the area around cilia (∼0.7 μm).

D-cells are much narrower, almost square in cross-section, shorter in proximal-distal direction. Their frontal side also bears multiple cilia which constitute the second, abfrontal, half of the lateral ciliary row. The basal complex of these cilia includes a single basal body, a basal foot and two rootlets, which are arranged identically to those in C-cells (Figure 2f, cell on the right). The axial rootlet also points basally and abfrontally, while the lateral rootlet runs more sharply downwards and frontally.

E-cells never bear cilia. They are somewhat flattened in shape and extend very short cytoplasmic flaps over the abfrontal sides of the adjacent D-cells. Together with intermittent F-cells they cover the entire abfrontal surface of the tentacle. At least some of these cells are ciliated. The ciliary basal complex may include a single kinetosome anchored to the apical cell membrane with short microtubules and bearing a cross-striated axial rootlet which runs diagonally in proximal-basal direction. Sets of two kinetosomes, arranged at right angles, and cupped by a Golgi apparatus, were also found (Figure 2g). Owing to the low frequency of F-cell cilia, we were unable to establish a more detailed model of the basal complex or verify if the basal feet are present or missing.

On the apical surface, all cells bear microvilli and cuticle. In all four species microvilli are longest on the frontal surface of the tentacle and near the lateral cilia (Figure 2b). The cuticle is two or three-layered. If present, the top layer is an indistinct felt-like structure composed of fine electron-dense strands attached to the tips of the microvilli. The medial layer comprises a thin horizontal skein of electron-dense strands radiating subterminally from the microvilli tips. The bottom layer is homogeneous and somewhat electron-lucent. The last two layers may be iterated two or more times along the length of the microvilli (Figure 2d,e).

Cell contacts are typical for bryozoans and include belt desmosomes in the apical position and septate junctions below (Figure 2f). The basal and lateral cell surfaces do not form interdigitating folds (unlike in phylactolaemate bryozoans).

Underneath the outer epithelium we found six basiepithelial nerves: one frontal, two lateral– frontal, two abfrontal–lateral and one abfrontal (Figure 2h).

A tube of relatively thick (∼0.3–0.8 µm) ECM provides axial support to the tentacle. On the inside it is lined by coelothelium and contains a coelom, at times occluded by somata of peritoneal cells. As in other bryozoans, the frontal and abfrontal coelothelial cells are myoepithelial, with contractile portions in contact with the ECM and anchored to it by hemidesmosomes. In some of the tentacles (only 1—2 per lophophore) we also saw a small muscle running basiepithelialy on the abfrontal surface of the tentacle (Figure 2b, tinted red). These muscles occurred in all studied species and had no relation to the distance to the tentacle base.

Two lateral coelothelial cells lack ciliation or contractile bundles, but in the distal half of the tentacle we often found well-developed secretory apparatus in the form of numerous, long cisternae of rER arranged in loose circles around the nucleus. We were unable to find conclusive proof of the existence of the sub-and epiperitoneal cells on the lateral sides of the tentacle coelom.

### 3.2 Lophophore base and the mouth region

The individual tentacles merge at the base of the lophophore (Figures 3, 4): at first the epidermal cells come into contact and the respective tentacle ECMs join more proximally. In both cases the abfrontal-lateral portions merge first, while frontal parts remain distinct for longer (Figure 3b). The fate of the frontal and abfrontal structures of the tentacles also differ markedly. The frontal cells transition into the mouth epidermis, and the frontal ECM forms the roof of the ring canal (Figures 3a, 4). The abfrontal cells make the outer lining of the lophophore base, and the abfrontal basement membrane becomes a thick ECM cylinder—the greatest non-skeletal support element of the polypide—and later forms the floor of the ring canal (Figures 3, 4).

**Figure 3.**
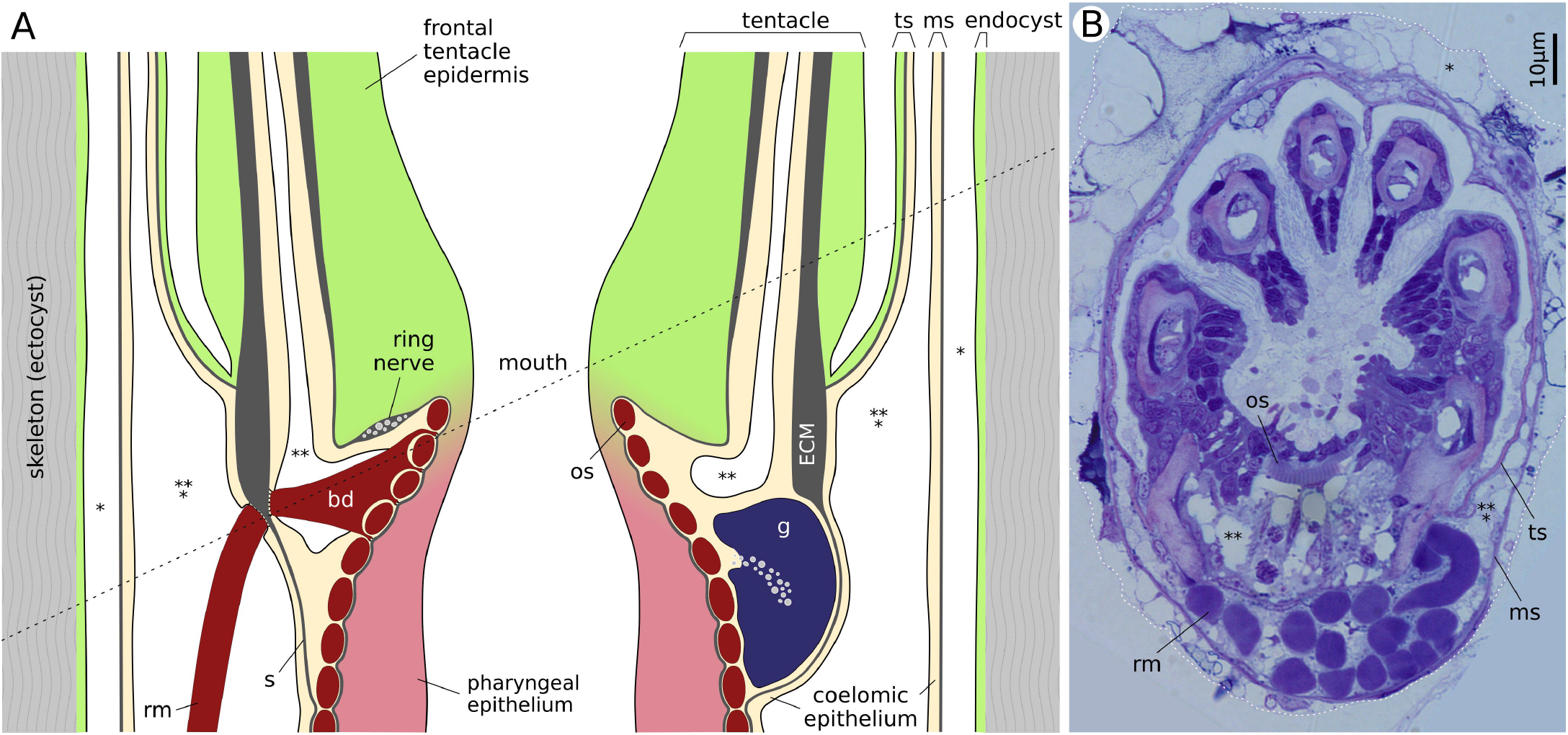
Lophophore base and mouth region. (a) Schematic drawing of a sagittal section through the lophophore base. Black dotted line shows the approximate position of the section in (b). (b) Obliquely transverse section through the zooid tube (white dotted line) and the polypide at the level of the mouth (*H*. sp. 2). ECM, extracellular matrix; g, ganglion; ms, membranous sac; os, oral sphincter, continuous with circular muscles of the pharynx; rm, retractor muscle; s, septum; ts, tentacle sheath; *, exosaccal cavity; **, lophophore coelom; ***, endosaccal cavity = trunk coelom

**Figure 4.**
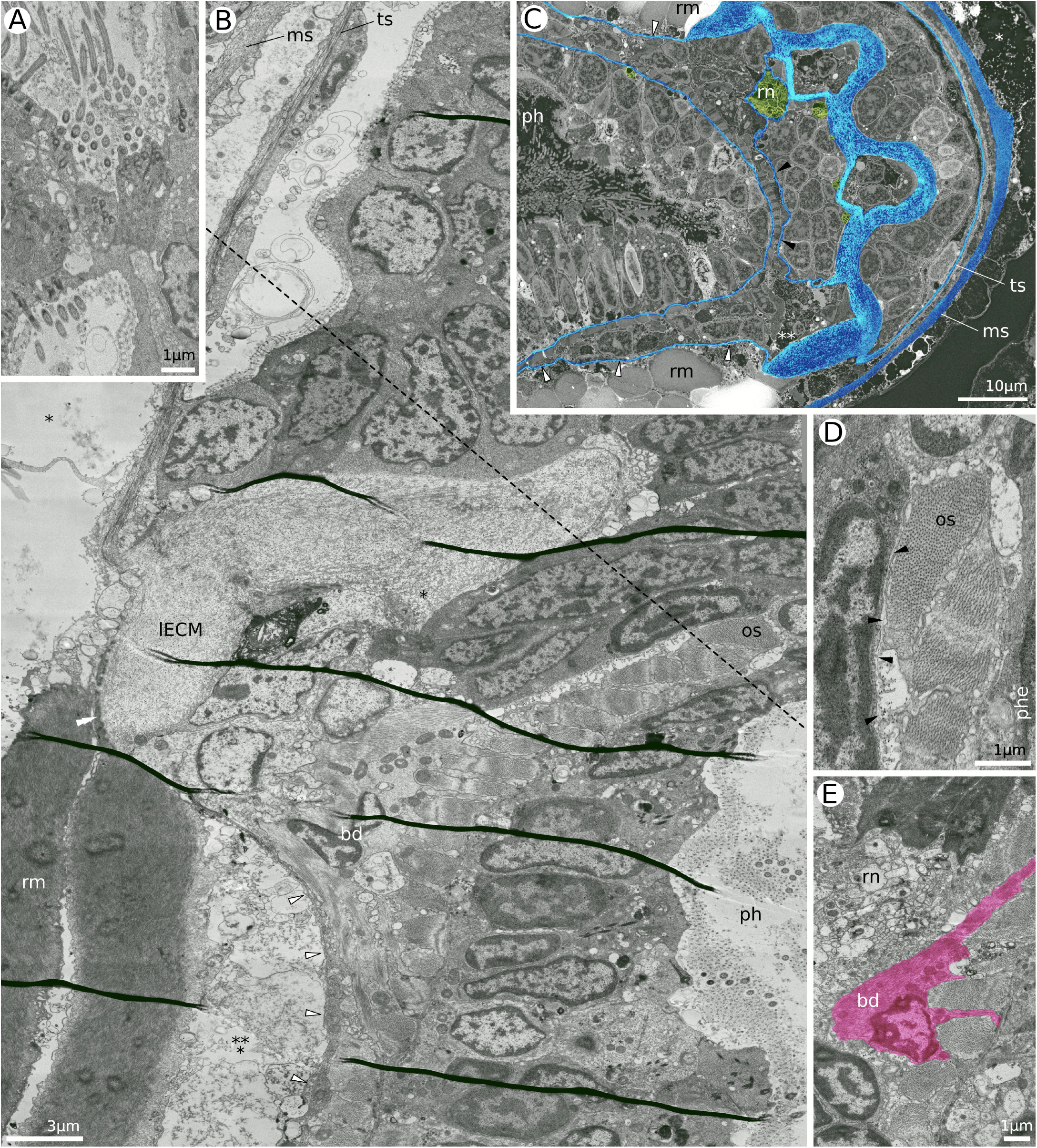
Lophophore base and mouth region. (a) Cross-section of merging tentacles of a protruded polypide (*H*. sp. 2, frontal/oral side up). (b) Midsagittal section through the mouth and lophophore base of a retracted polypide showing the origin of the tentacle sheath, the thick ECM collar at the lophophore base, oral and pharyngeal sphincter and lophophore–trunk septum (*H. robusta*). Dotted line represents the approximate location of section (c). (c) Obliquely transverse section through the tentacle base and mouth region (*H. robusta*, all ECM blue, nerves green). (d) Cross-section through the oral sphincter showing ECM of the lophophore roof folding over the muscles and continuing onto the pharyngeal wall (*H. robusta*). (e) Sagittal section through the mouth showing ring nerve and buccal dilator (*H. robusta*). bd, buccal dilator; lECM, extracellular matrix at the base of the lophophore; ms, membranous sac; os, oral sphincter, continuous with circular muscles of the pharynx; ph, pharynx; phe, pharyngeal epithelium; rm, retractor muscle; rn, ring nerve; ts, tentacle sheath; *, exosaccal cavity; **, lophophore coelom; ***, endosaccal cavity; black arrowhead, ECM of the lophophore roof; white arrowhead, lophophore–trunk septum; double white arrowhead, lophophoral insertion of retractor muscles

Above the mouth, the A-, B-and C-cell lines continue onto the preoral region, whereas D-, E-and F-cells appear on the outer surface of the lophophore. As a result, lateral ciliary rows split into frontal and abfrontal portions (Figure 4a). As the C-cells of neighboring tentacles come into contact, the density of ciliary cover decreases and ciliation gradually disappears. Proximally to the last protruding axonemes there are several irregularly arranged naked kinetosomes. Abfrontally, D-cells meet while still bearing cilia, but the ciliation ends quickly. No trace of ciliated pits, described in gymnolaemate bryozoans, was seen, although we observed mitoses at the tentacle bases.

The ECM near the tentacle base develops thick abfrontal–lateral horns (up to 10 times thicker than the frontal ECM) which elongate and merge just above the mouth (Figure 4b,c). The basement membranes of the frontal tentacle surfaces retain individuality and continue proximally below the level of the oral sphincter for 5 to 10 μm before merging (Figure 4c). After joining together, they abruptly become very delicate and hard to trace even on TEM images. As the tentacle coeloms open up into the ring canal of the lophophore, the unified preoral ECM slopes up towards the oral sphincter, folds over the sphincter muscles and continues downwards as the basement membrane of the pharyngeal epithelium (Figure 4b,d).

On the abfrontal side, the outer ECM remains thickened for some distance below the mouth, where it supports a number of powerful muscles and also gives rise to the tentacle sheath (Figure 4b). The latter merges with the lophophore ECM slightly below the mouth. The thick outer ECM of the lophophore continues proximally for an additional 10–15 μm (i.e. for ∼25 μm below the oral sphincter). Large buccal dilator muscles, each of which is an individual cell, cross the ring canal (Figure 4b,e). As they pass between the circular muscles of the oral sphincter, buccal dilators attach to the ECM of the mouth in several places and run radially towards the thick outer ECM. Opposite the buccal dilators, in the endosaccal cavity of the trunk, the outer ECM cylinder also supports the retractor muscles, which attach to its proximal surface around most of the lophophore perimeter (Figures 3b, 4b,c).

Below the attachments of the buccal dilators and retractors, the lophophoral ECM abruptly becomes very delicate and slopes down towards the pharynx or around the ganglion (Figure 4b,c). This delicate extracellular layer forms the floor of the ring canal, i.e. the septum separating lophophore and trunk coeloms. The ECM of the septum does not simply merge with the ECM of the pharynx wall. Instead, it forms an outer “leaf” going over the circular muscles of the pharynx (Figure 4c).

The circumoral nerve ring is located in the lophophore coelom, in association with the ECM (Figure 4e). It is, however, impossible to tell exactly how these two structures correspond, since the ECM is so delicate and hard to trace.

The ganglion is a relatively small, compact structure sitting between the pharynx and the terminal part of the rectum (Figure 5a). The neuropile forms an invagination in the distal half of the ganglion pointing diagonally towards the mouth (Figure 5b). The distal boundary of the ganglion coincides with the lower parts of the buccal dilators (Figure 5a,c), so that the top of the ganglion is bracketed by two muscle cells. More proximally the ganglion acquires two lateral lobes, similar in appearance to lateral ganglia in *Cinctipora elegans* (Schwaha et al., 2018). The lobes are compact elongated structures made of cells with electron-dense cytoplasm (Figure 5a). We detected no neuropile and have no immunoreactivity data on these structures, thus we hesitate to call them lateral ganglia.

**Figure 5.**
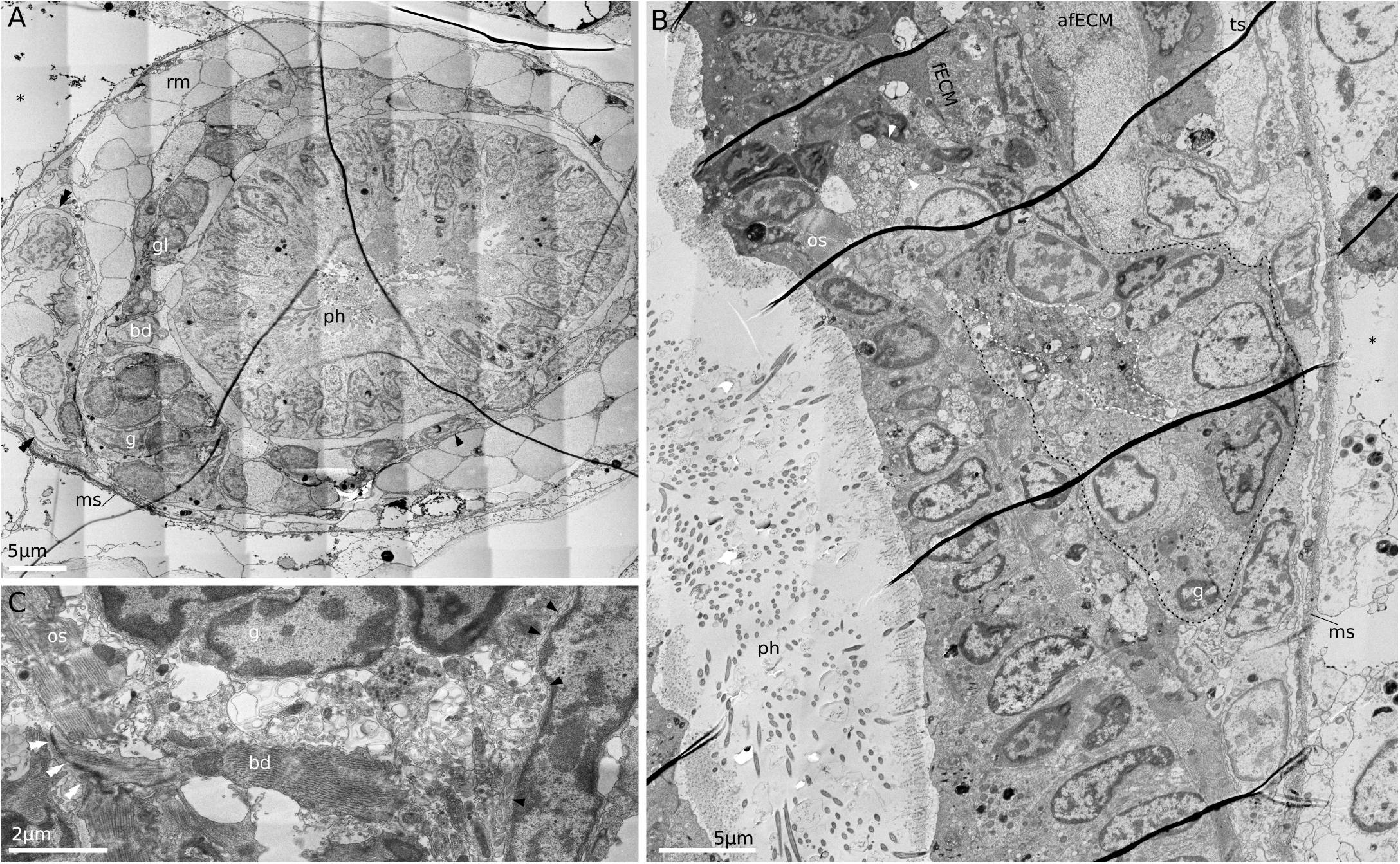
Ganglion. (a) Cross-section through the zooid tube with retracted polypide at the level of the ganglion, showing the distal portions of the ganglion lobes (Hornerid gen. sp3). (b) Midsagittal section through the ganglion of a retracted polypide (*H. robusta*, ganglion surrounded by black dotted line, neuropile—by white dotted line), (c) Cross-section of a retracted polypide through the ganglion, showing buccal dilator, oral sphincter, lophophore–trunk septum and neuropile (*H*. sp. 2). afECM, abfrontal tentacle ECM; bd, buccal dilator; fECM, frontal tentacle ECM; g, ganglion; gl, ganglion lobe; ms, membranous sac; rm, retractor muscle; ts, tentacle sheath; os, oral sphincter; * - exosaccal cavity; black arrowhead, ECM of the septum; white arrowhead, ring nerve; black double arrowhead, distal part of the rectum; white double arrowhead, hemidesmosomes attaching buccal dilator to the oral ECM

The tentacle sheath is a cylindrical part of the polypide body wall stretching from the origin at the base of the lophophore to the end point where it merges with the vestibular wall and the membranous sac (Figure 6a, white dotted line). In the retracted polypide the tentacle sheath forms the casing around the tentacles, creating a temporary atrial cavity within. When the polypide is protruded, it forms a flexible stalk that supports the tentacle crown. The location of the point where the tentacle sheath transitions into the membranous sac and the vestibular wall (Figure 6A) is relatively stable regardless of the position of the polypide. It is located approximately 100–150 μm distal of the aperture (which corresponds to the cardia–caecum boundary in a protruded polypide). The origin of the tentacle sheath, which is located at the lophophore base (Figure 6b), moves up and down with the lophophore.

**Figure 6.**
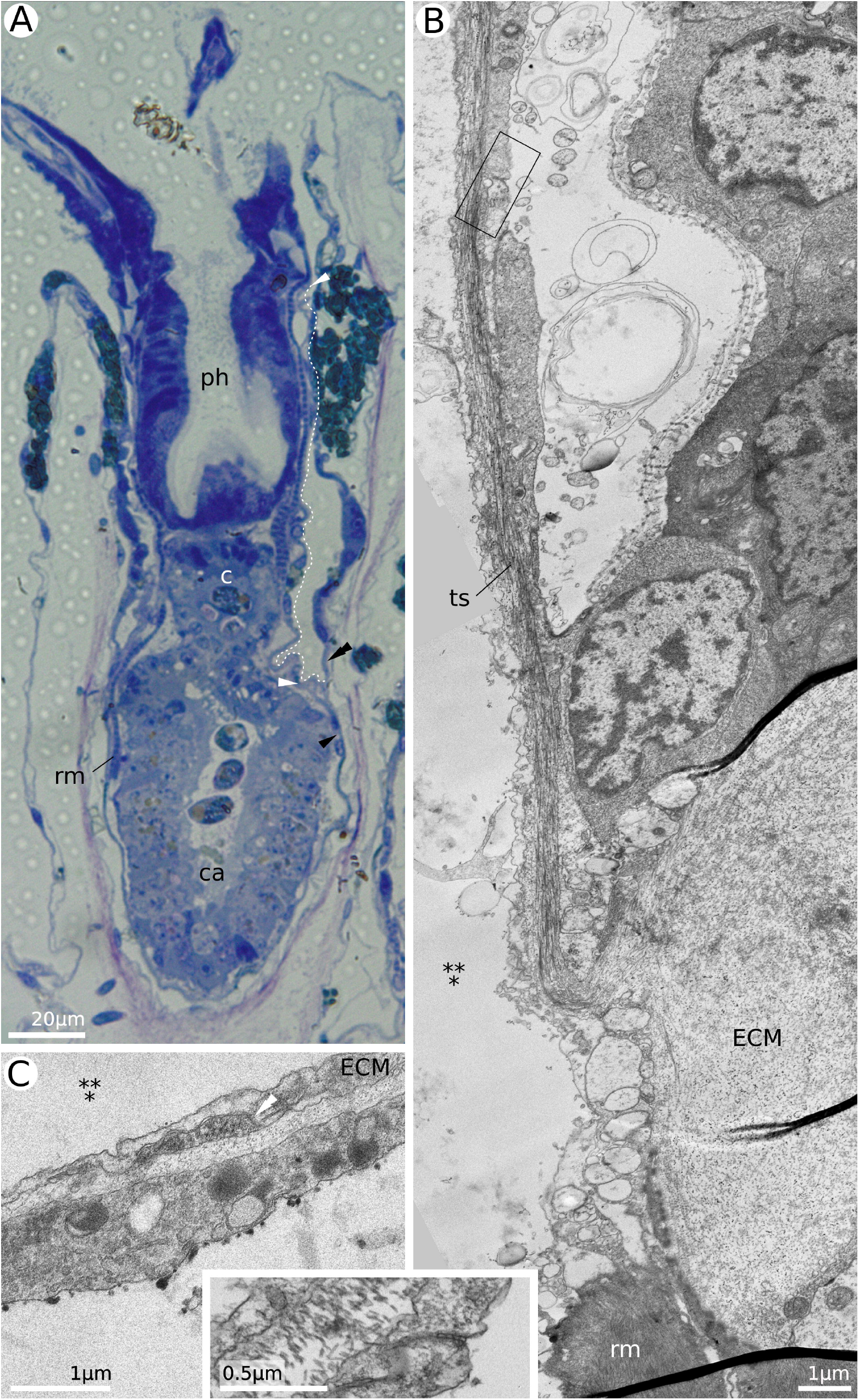
Tentacle sheath. (a) Obliquely frontal section through the protruded polypide (*H*. sp. 2). (b) Longitudinal section through the base of the tentacle sheath in a retracted polypide, showing lack of cuticle on the tentacle sheath epidermis (*H. robusta*). Inset (*H*. sp. 1) shows septate cell junction similar to the black frame in figure (b). (c) Transverse section through the tentacle sheath of a protruded polypide showing longitudinal muscle (*H*. sp. 2 endosaccal cavity above, vestibular space below). c, cardia; ca, caecum; ECM, extracellular matrix; ph, pharynx; rm, retractor muscle; ts, tentacle sheath; ***, endosaccal cavity

The tentacle sheath is a fine structure ∼1–2 µm thick with squamous coelothelial and epidermal cells resting on a 15–45 nm ECM. There is no cuticle. Tentacle sheath musculature is represented by very delicate longitudinal muscles (10–15 fibers per cell; Figure 6c). Although the surface is not ciliated, we found a free kinetosome in one of the epidermal cells.

The epidermis of the tentacle sheath continues onto the vestibular wall, whereas the coelothelium goes onto the inside of the membranous sac. The ECM material is equally thick in all three structures. More details on the composition of the membranous sac, vestibular wall and terminal membrane will be given in our following publication.

### 3.3 Retractor muscles

There are about 50 individual retractor muscle cells in a zooid. They originate from paired semicircular attachment zones on the abanal surface of the zooid wall, about 400 µm from the aperture, and connect the polypide to the skeletal wall (Figure 7a,b). In agreement with the protrusion distance of the polypide, the retractor muscles undergo substantial changes in length: from ∼50 µm when contracted to ∼370 µm when relaxed (Figure 7a).

**Figure 7.**
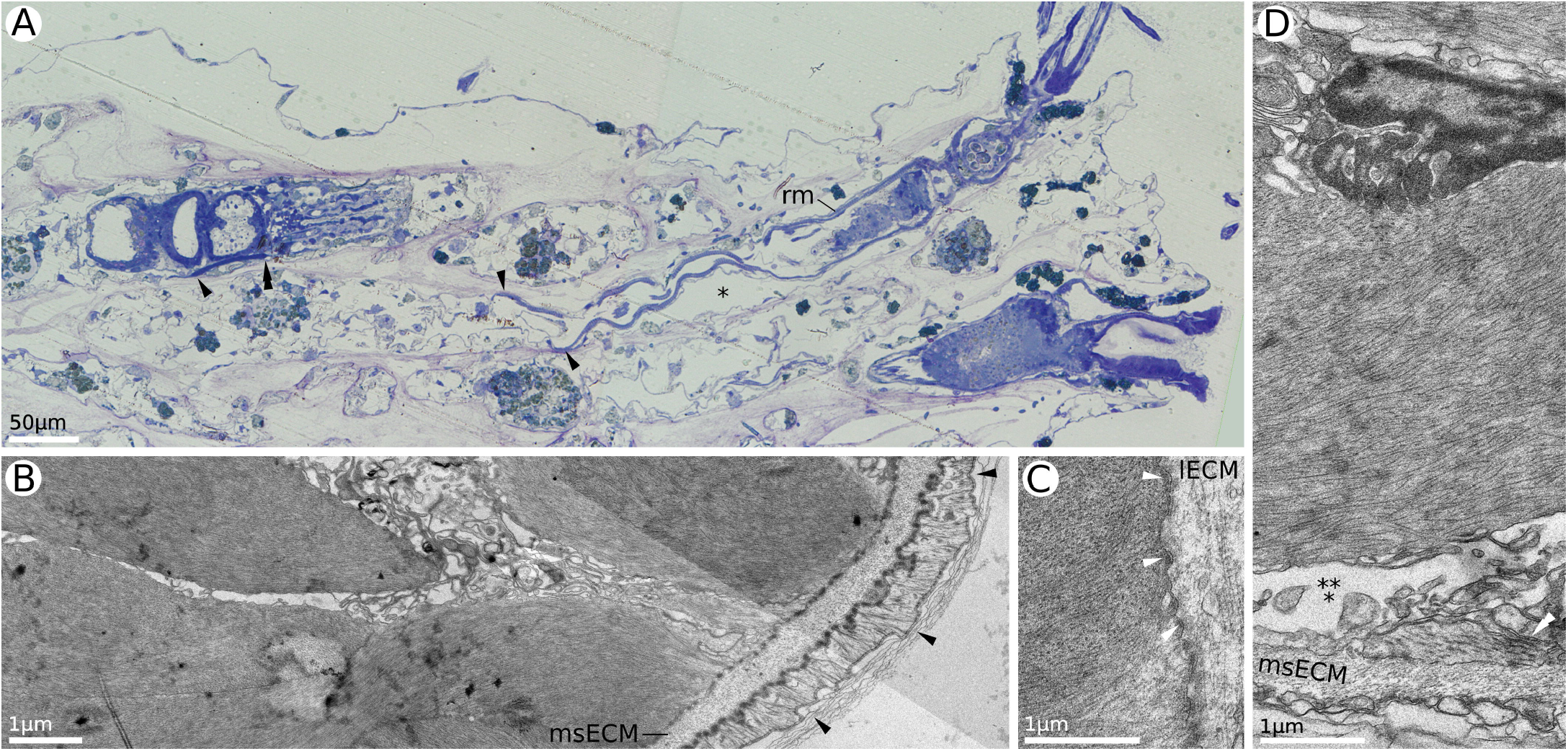
Retractor muscles. (a) Sagittal section through colony branch showing two expanded (distal) and one retracted polypide (proximal), illustrating retractor muscles in the relaxed and contracted state respectively (*H*. sp. 2). (b) Transverse section through the pharyngeal region of the retracted polypide showing skeletal origins of retractor muscles (*H*. sp. 2). (c) Details of lophophoral insertion of the retractor muscle showing hemidesmosomes (*H*. sp. 1). (d) Cross-section through the medial part of the contracted retractor muscle showing nuclear indentations (*H*. sp. 2). lECM, lophophoral extracellular matrix; msECM, extracellular matrix of the membranous sac; rm, retractor muscle; *, exosaccal cavity; ***, endosaccal cavity; black arrowhead, skeletal attachments of retractor muscles; double black arrowhead, lophophoral insertion of retractor muscles; white arrowhead, hemidesmosomes at lophophoral insertion; double white arrowhead, membranous sac musculature

The muscles extend proximally towards the basal parts of the lophophore ECM (Figures 4b, 7c). In retracted polypides they are arranged in a semicircle cupping the abanal side of the downward branch of the gut, or concentrate in two groups in the space between gut branches. More proximally, retractor muscles spread out to a near-complete circumference, and the lophophore-insertion area encloses most of the lophophore, except for a gap around the ganglion (Figure 5a).

Each retractor muscle is an individual cell with unusually shaped nucleus which forms numerous interdigitating protrusions at the interface with the contractile bundles (Figure 7d). Retractor muscles are enclosed within the membranous sac for their entire length. Both at the origin and at the lophophore insertion they are anchored to the ECM with hemidesmosomes (Figure 7c).

### 3.4 Digestive system

The digestive system of hornerids has a typical bryozoan composition: a downward branch (mouth, pharynx and cardia: Figure 8), a blind sac-like caecum, and an upward branch (pylorus, rectum and anus: Figure 9). The downward branch usually follows the abfrontal surface of the zooid tube and upward branch runs along the frontal surface.

**Figure 8.**
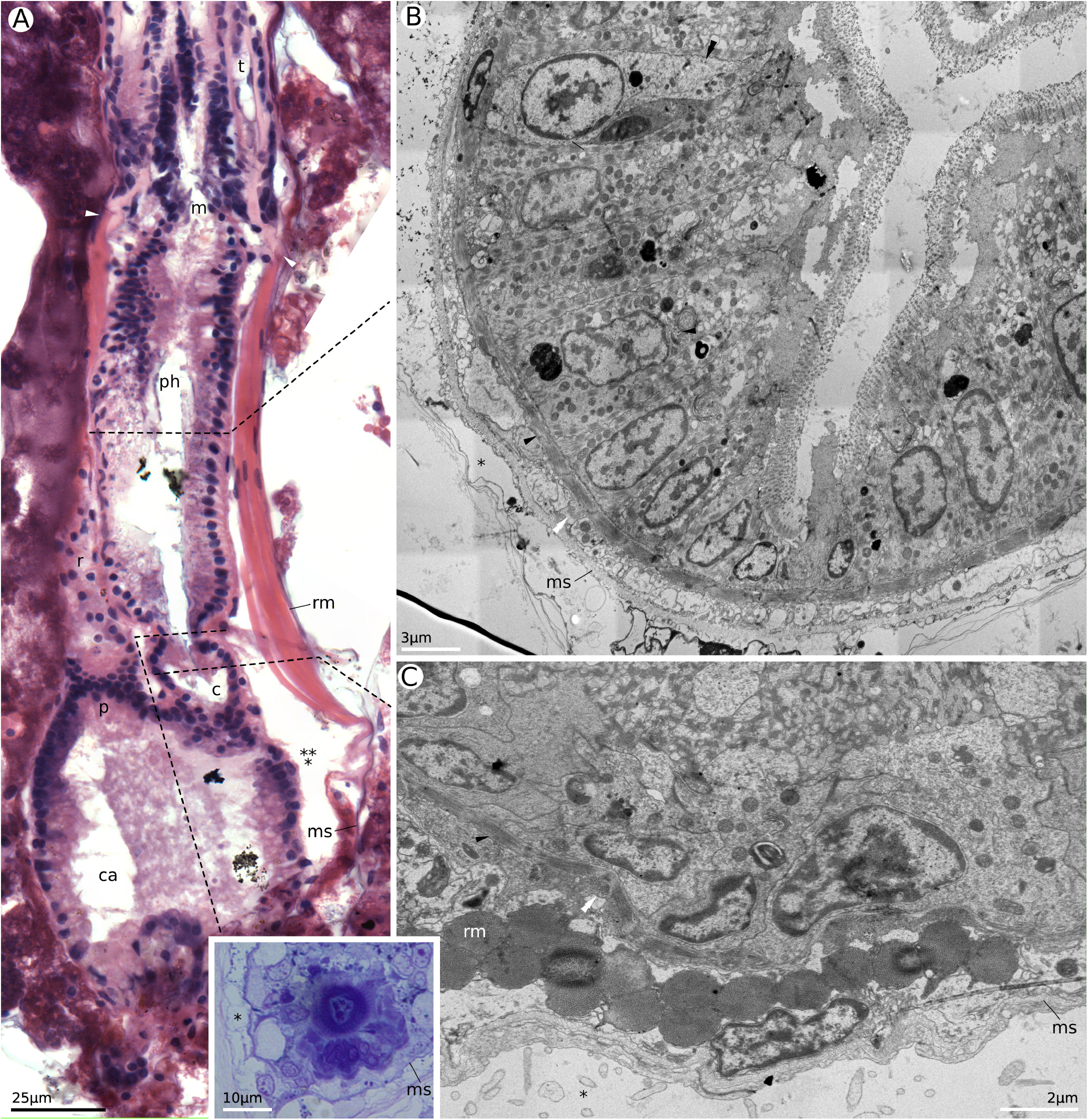
Digestive system: downward branch. (a) Oblique sagittal section through the partially retracted polypide showing digestive system. Black dotted lines indicate approximate locations of the cross-sections (*H. robusta*). (b) Cross-section through the medial part of the pharynx of a retracted polypide showing myofibrils in the pharyngeal epithelium, circular and longitudinal muscles of the pharynx wall (*H. robusta*). (c) Cross-section through the cardia of a protruded polypide (*H*. sp. 2); inset shows cross-section of a contracted pharyngeal–cardial valve (*H*. sp. 1). c, cardia; ca, caecum; m, mouth; ms, membranous sac; p, pylorus; ph, pharynx; r, rectum; rm, retractor muscle; t, tentacle; *, exosaccal cavity; ***, endosaccal cavity; black arrowhead, circular musculature of pharynx and cardia; double black arrowhead, non-muscular cell in pharyngeal epithelium; white arrowhead, lophophoral insertion of retractor muscles; double white arrowhead, longitudinal muscles of pharynx and cardia

**Figure 9.**
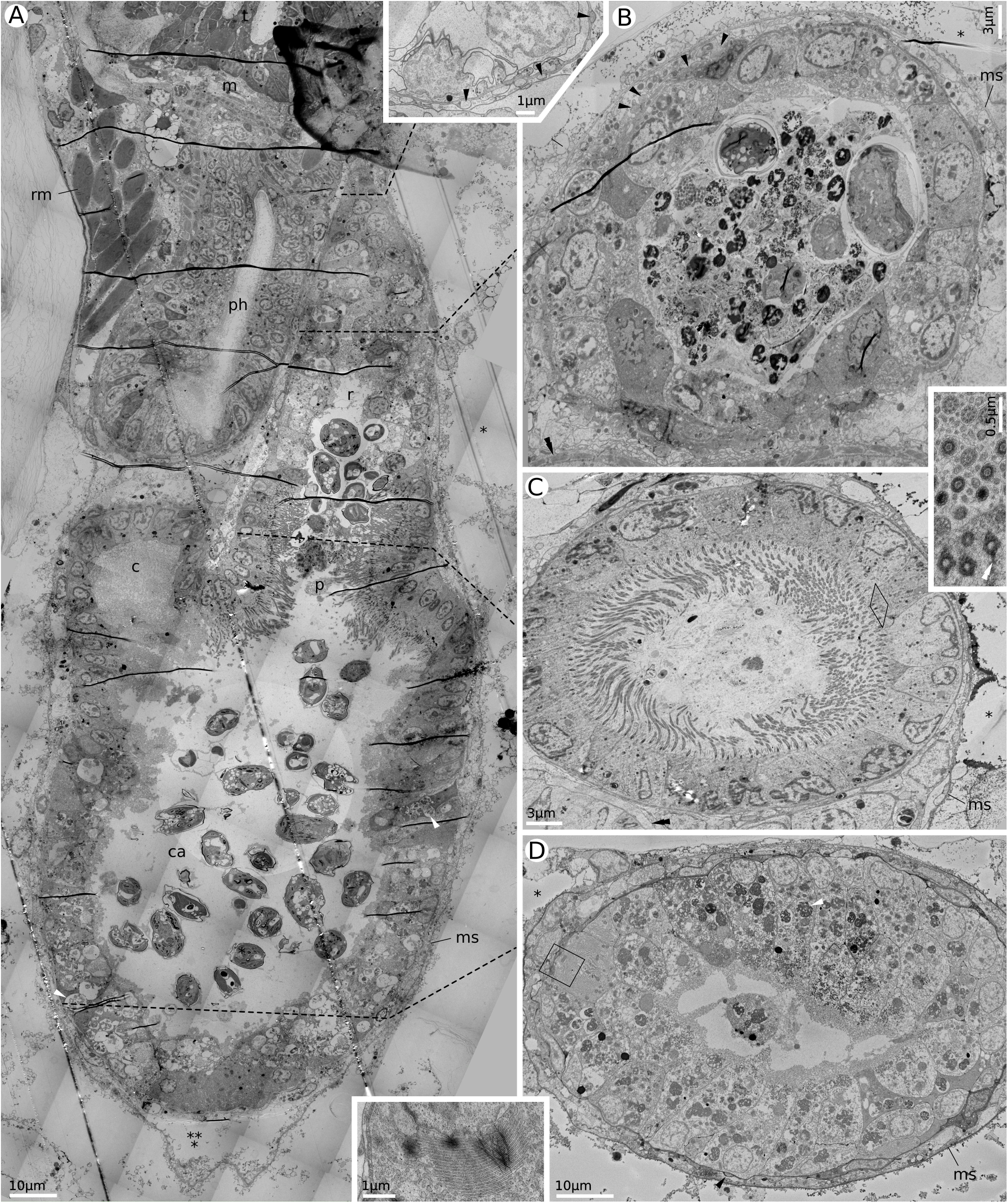
Digestive system: upward branch. (a) Sagittal section through retracted polypide showing digestive system. Black dotted lines indicate approximate locations of the cross-sections (*H. robusta*). Inset presents rectum near anus (Horneridae gen. sp. 3). (b) Cross-section of the rectum with a fecal pellet inside (*H. robusta*). (c) Cross-section of the pylorus (Horneridae gen. sp. 3). Black frame corresponds to the tilted area, similar to the inset. Inset shows details of pyloric cilia and microvilli (*H*. sp. 1). (d) Cross-section through the proximal part of caecum, showing active biosynthetic apparatus and large secondary lysosomes (*H. robusta*). Black frame corresponds to the area similar to the inset. Inset demonstrates numerous rER cisternae (Horneridae gen. sp. 3). c, cardia; ca, caecum; m, mouth: ms, membranous sac; p, pylorus; ph, pharynx; r, rectum; rm, retractor muscle; t, tentacle; *, exosaccal cavity; ***, endosaccal cavity; black arrowhead, longitudinal gut musculature; double black arrowhead, circular muscles of the pharynx; white arrowhead, secondary lysosomes; double white arrowhead, basal feet of pyloric cilia

The digestive system starts with a round muscular mouth. During normal feeding the mouth is kept open at ∼25 µm in diameter (it can open even wider for swallowing), while in fixed material it is usually more contracted at ∼13 μm in diameter. The circular cross-striated muscles around the mouth make up the oral sphincter. Interestingly, the oral sphincter can be identified in living polypides by differential contraction, but has no boundary from the pharyngeal muscles below (Figures 4b, 5b). The epithelium of the mouth itself is separated clearly into the upper and lower parts (Figure 5b). The upper part is made of epidermal cells continuous with the frontal cells of the tentacles. They typically have electron-dark cytoplasm and no ciliation. The lower portion of the mouth is lined with occasionally ciliated myoepithelium with distinctly more electron-lucent cytoplasm and an interior composition similar to pharyngeal cells. Thus, the mouth has a more functional than anatomical identity.

The pharynx is a direct continuation of the mouth. The outer contours of the pharynx are oval in cross-section, but the lumen is Y-shaped (Figure 8b). The latter form arises from the uneven height of the cells (from ∼5 μm between ridges to ∼15 μm on the ridge). All cells are myoepithelial, with the contractile portions arranged vertically, i.e. in apical–basal direction, around the nucleus (Figure 8b). We saw no structural differences in the cells in and between ridges and no indication of anchor cells *sensu* Nielsen (2013). Their apical surfaces bear long (∼1 μm) and dense microvilli. The pharynx is the only region of the gut where we observed cell divisions.

A cuticle layer continues from the tentacle surface over the mouth and onto the pharynx, and retains its three-layered structure. An upper layer, thick and felt-like, comprises fine electron-dense strands (antennulae) radiating upwards from the tips of the microvilli. The medial layer comprises a flat skein of electron-dense strands radiating horizontally from the microvilli halfway along their length (cf. the analogous but subterminally located layer on the tentacles). The third, bottom layer is fine-grained, homogeneous and electron-lucent. Cell cytoplasm contains an elongated nucleus, mitochondria and rER, but no major vacuoles.

In the distal half of the pharynx, pharyngeal ridges can bear ciliation. In *H. robusta, H*. sp. 2 and Horneridae gen. sp. 3 this comprises 5–15 multiciliated cells located approximately in the center of each ridge. Each cilium has a single kinetosome, axial and lateral rootlets and a proximally directed basal foot. In *H*. sp. 1 the number is smaller: from 1 to 7 cells. We were unable to positively confirm that they are monociliated, but the basal apparatus includes two kinetosomes instead of one, which implies monociliated condition.

In the proximal portions cilia are missing in all of the species. The ridges tend to be higher and the pharyngeal lumen smaller.

The pharynx is supplied with basiepithelial nerves. In *H. robusta* there are two or three major pharyngeal nerves, each located underneath the ridges. In *H*. sp. 2 we found a large and variable number of nerves. In addition, this species alone possessed large intercellular spaces, which can be mistaken for vacuoles in semi-thin sections.

Pharyngeal epithelium rests on a delicate ECM with embedded cross-striated circular muscles (Figure 8b). This layer of musculature is a seamless continuation of the oral sphincter. On the outside the pharynx is also supplied with irregular longitudinal muscles (10-20 fibers per cell; Figure 8b). In *H. robusta* and *H*. sp. 1 the number is low (∼5), whereas in *H*. sp. 2 these muscles are more numerous (∼20).

A very narrow constriction or valve, supplied with powerful circular muscles, separates the pharynx from a short cardia (Figure 8a, inset). The cardia is lined by cuboidal epithelial cells ∼4-5 µm tall (Figure 8c). Apical surfaces of these cells have no ciliation but microvilli are present. The circular and longitudinal musculature present in the pharynx continues onto the outer cardial wall (Figure 8c).

The caecum is the largest portion of the gut (Figures 8a, 9). Caecal cells are typically tall (∼10-20 µm) and bear no ciliation. In the distal part of the caecum the cells often have electron-dark cytoplasm, low-contrast nuclei and almost no inclusions. In the medial part of the caecum some cells engage in active biosynthesis as evidenced by enlarged perinuclear spaces and numerous rER cisternae, often enlarged or else forming densely packed fields and occupying nearly all the cytoplasm. More proximally, a different type of cell becomes dominant. These cells have electron-light cytoplasm, high-contrast nuclei and great numbers of large (up to 5 μm in diameter) organelles which we interpret as secondary lysosomes with a variety of electron-lucent and electron-dense contents (Figure 9a,d). Towards the proximal end of the caecum, the number of cells with secondary lysosomes grows, and the content of the latter becomes more granulated and electron-dark. On the sections examined by transmitted-light microscopy these organelles have a distinctive orange-brown coloration, indicating the presence of lipofuscin. On any given cross-section one may see up to 20 such vacuoles per cell. Nevertheless, cells with active rER are also present throughout the proximal part of the gut and particularly near the origin of the funiculus (Figure 9a,d, inset). At no point in the caecum did we observe cell divisions. The outer caecum wall lacks prominent circular musculature. Several diagonal and ∼30 delicate longitudinal muscles encase the caecum between the ECM and coelothelium.

The pylorus is a relatively short segment of the gut (∼10–20 cells in length; roughly equivalent to cardia), lined by cylindrical to cuboidal cells bearing dense ciliary and microvillar cover (Figure 9a,c). The basal complex of a pyloric cilium includes a very prominent axial rootlet, a basal foot and short lateral rootlet (Figure 9c, inset). The basal complexes are arranged somewhat irregularly, with basal feet of neighboring cilia pointing in different directions. On the outside the pylorus is surrounded by fine ECM, delicate longitudinal muscles, and coelothelium.

The pylorus transitions to the rectum without a valve (Figure 9a). The cells lining the rectum are relatively large and cuboidal (∼6 μm), usually without distinctive apical structures (Figure 9a,b). Some of these cells tend to have electron-lucent cytoplasm and large secondary lysosomes reminiscent of the caecum wall. Others show signs of active biosynthesis. Outer layers include the same components: ECM, longitudinal muscles, and coelothelium. Interestingly, no distinctive anal sphincter was seen in any of the studied species (Figure 9a, inset).

### 3.5 Funiculus and trunk coelom

The funiculus of the studied hornerids is a solid ECM cord ∼4 by 10 μm in cross-section, devoid of either lumen or lacunae (Figure 10). In a relaxed state it is ∼250 μm long, whereas contracted it is about 50–150 μm. Two funicular muscles are adpressed to the outside the ECM cord (Figure 10a,b). At their origin, each muscle cell is anchored to the membranous sac and — through it — to the cystid at the funicular attachment zone (Figure 10c). In addition, they are attached to the ECM cord of the funiculus itself with hemidesmosomes. The ECM of the funiculus joins seamlessly with the ECM of the outer gut wall (Figure 10b) as well as the membranous sac (Figure 10c). The funicular attachment footprint is 40–50 μm in proximal-distal direction and ∼30 μm laterally. We detected no mural pores on the cystid wall at the point of attachment, which suggests that the funiculus does not take part in interzooidal transport or neural connectivity.

**Figure 10.**
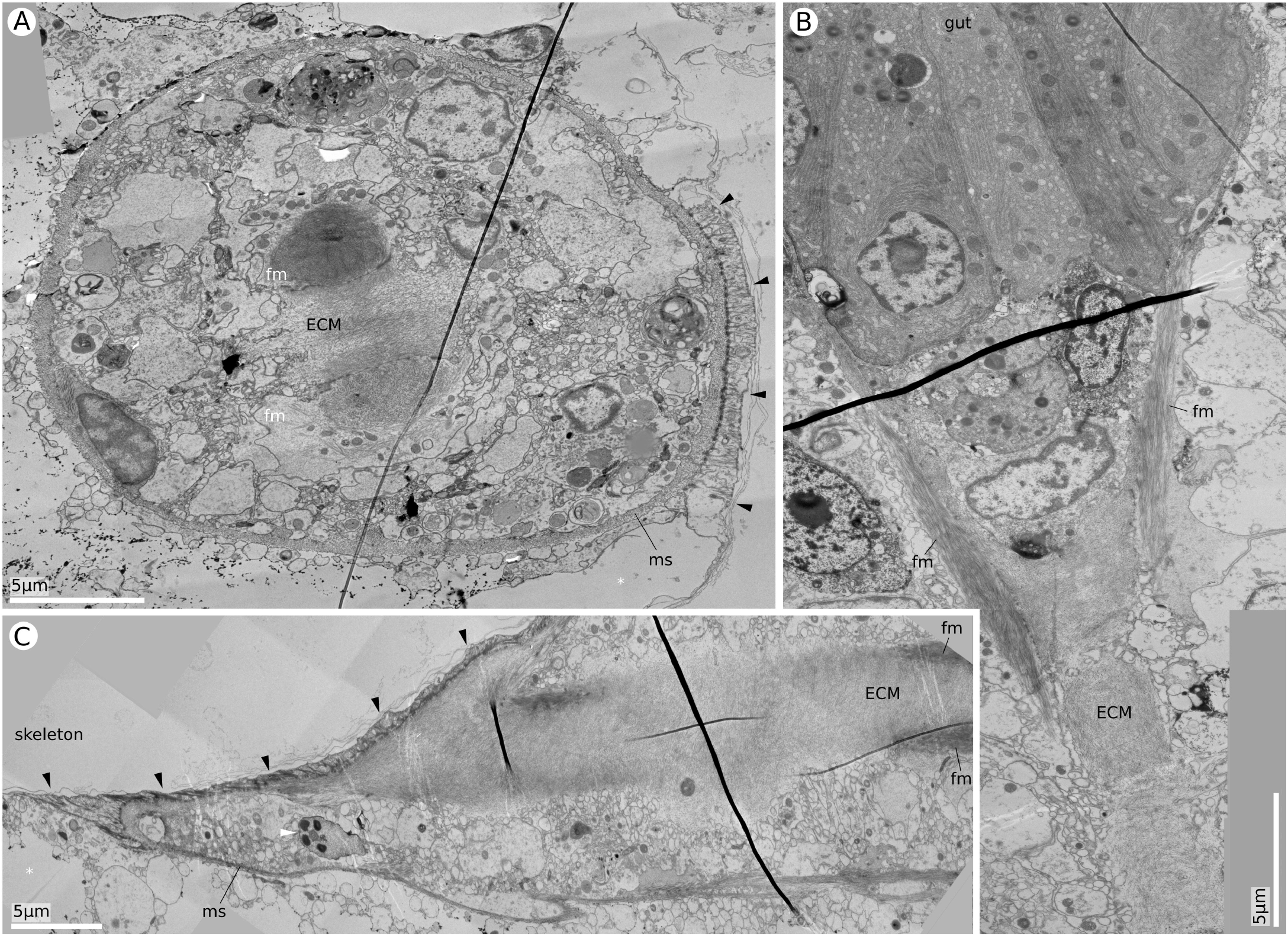
Ultrastructure of the funiculus. (a) Cross-section through the zooidal chamber at the proximal end of the membranous sac and funiculus, showing the distalmost edge of the funicular attachment ligament (black arrowheads) (*H*. sp. 2). (b) Longitudinal section through the proximal tip of the gut and the origin of funiculus (*H. robusta*). (c) Longitudinal section through the funicular attachment (*H. robusta*, proximal direction on the left). ECM, extracellular matrix; fm, funicular muscle; ms, membranous sac; black arrowhead, tendon cell of the funicular attachment; white arrowhead, sperm cell; *, exosaccal cavity

At the base of the funiculus we sometimes saw a group of cells dissimilar from gut cells (Figure 10b). We did not see specific insertion points of the funicular muscles into the gut wall, so it is possible that they continue along the outer surface of the gut as longitudinal caecum muscles (as in *Crisia eburnea*, Worsaae et al., 2018). If so, they cannot be distinguished from other caecum musculature.

The trunk coelom (endosaccal cavity) generally contains numerous coelomocytes, especially noticeable in the distalmost portion of the membranous sac of retracted polypides (Figure 10a). In addition to cells, we also noted numerous membrane-bounded vesicles (potential cell debris).

## 4. Discussion

Polypides of the four studied hornerid species have a relatively typical anatomy compared to other cyclostomates. Except for the funiculus, the broad structure of the organs and body compartmentalization, as well as some ultrastructural details, are in agreement with studies on other species and often show clear functional linkages.

Gross tentacle structure of our studied species is in line with all other examined cyclostomates, although, like other members of this order, hornerids lack ciliated pits, recently explored in gymnolaemates (Shunatova & Borisenko, 2020). The general principles of organization of the basal apparatus of lateral-frontal and lateral cilia are relatively conservative in Bryozoa. Lateral-frontal cilia always have two kinetosomes and a combination of axial and diagonal rootlets, arranged similarly relative to the tentacle midline, which implies shared and conserved function (both motile and mechanoreceptive). The differences in basal complexes of *Hornera*, phylactolaemates and gymolaemates are slight. In Phylactolaemata there is an additional diagonal rootlet, absent in our studied species, whereas freshwater species lack a basal foot which is present in hornerids. Images of the lateral–frontal cilia in Gymnolaemata by Lutaud (1973) show one axial rootlet proceeding from the main kinetosome and going through the groove in the nucleus. An additional kinetosome is displaced toward the tentacle midline. There is a second axial rootlet, arising from this kinetosome and joining the main axial rootlet, an arrangement that is similar to what we have found in hornerids. Finally, a third, diagonal rootlet attaches to the additional kinetosome. Shunatova (2002) also gives excellent descriptions of the basal apparatus of B-cells in *Rhamphostomella ovata* and *Eucratea loricata*. The main kinetosome gives rise to one axial rootlet and a basal foot pointing frontally. The additional kinetosome is also located frontally and gives rise to one axial and two diagonal rootlets (going proximally and distally along the tentacle). In addition, there is a microfilament bundle linking the kinetosomes.

Orientation of the basal feet corresponds to the direction of the active stroke of the motile cilia (e.g., Boisvieux-Ulrich et al., 1985). Although lateral-frontal cilia do not take part in generating water currents, they make occasional flicks in order to intercept and transport particles back into the center of the feeding current. The location of the basal foot, if present, is in agreement with this view, since it is always pointing frontally, towards the midline of the tentacle.

Organization of the lateral cilia in hornerids resembles that of gymnolaemates (Gordon, 1974; Lutaud, 1973; Smith, 1973). Lacking, however, are some of the cytoplasmic anchoring mechanisms found in phylactolaemates, i.e. one of the lateral rootlets and a network of microtubules radiating from the basal feet of lateral cilia (Fawcett 1958, Tamberg and Shunatova 2017). The presence and orientation of the basal feet, however, is universal in all bryozoans studied so far.

Abfrontal cilia are also found in all bryozoans and are widely considered to be mechanoreceptive and non-motile. Shunatova and Nielsen (2002) reported only monociliated cells on the abfrontal surfaces of the tentacles in four stenolaemate species (*Crisia eburnea, Crisiella producta, Tubulipora flabellaris* and *Patinella verrucaria*). The same authors reported that monociliated cells alternate with multiciliated ones which bear 7–15 non-motile cilia in 17 gymnolaemate species. Gordon (1974) also described both single cilia and tufts of ten cilia in the gymnolaemate *Cryptosula pallasiana*. In *Rhamphostomella ovata* a basal foot and two rootlets are reported (Shunatova and Nielsen 2002). In phylactolaemates the abfrontal ciliary arrangements are always made of paired monociliated and/or biciliated cells (Tamberg and Shunatova, 2017), giving rise to tufts of three or four cilia (Riisgård et al. 2004, Tamberg and Shunatova, 2017).

Ciliated mechanoreceptor cells are known to depolarize when the sensory cilia are displaced in the direction of the basal foot. In other words, the foot projects from the kinetosome in the direction of ciliary movement which excites the cell (Barber and Boyde, 1968). If the abfrontal cilia are indeed mechanoreceptive, we can make inferences about the ciliary displacements they are designed to detect. In Phylactolaemata both upper and lower cells of the abfrontal pair have distal-facing feet, i.e. they preferentially detect being displaced towards the tip of the tentacle. Unfortunately, in Gymnolaemata the direction of the basal feet of the abfrontal cilia has not been described, and in this study, we have not seen basal feet.

Although organization of the ciliary basal complexes differs somewhat in cyclostomates, phylactolaemates and gymnolaemates, all these groups share deep structural similarities.

Tentacle and lophophoral coeloms are completely separated from the trunk by a delicate unbroken septum. This observation agrees with previous reports on rectangulates, articulates and tubuliporines (Shunatova & Tamberg, 2019).

The tentacle sheath of all bryozoans is supplied with longitudinal and sometimes circular musculature. The role of these muscles seems to reside primarily in the execution of behavioral reactions, such as scanning (presumably locating a preferred feeding position), optimal positioning of the lophophores around the chimney, colonial cleaning and others (see Shunatova and Ostrovsky 2001, 2002 and references therein). In cheilostomatids the everted tentacle sheath is fully revealed above the skeleton, and polypides can perform a variety of inclines, rotations and bends (Winston 1978; Shunatova and Ostrovsky 2001, 2002). In cyclostomates, however, only the tentacles protrude from the aperture. With the tentacle sheath hidden in the skeletal tube, its movements are restricted. For instance, Shunatova and Ostrovsky (2001) describe no scanning in any of their studied cyclostomate species (*Crisia* sp., *Crisiella producta, Tubulipora flabellaris, Patinella verrucaria, Disporella hispida*). Our observations on living *H. robusta* and *H*. sp. 2 are in good agreement: the position of the lophophore may be adjusted, but only slightly, by minute bends and rotations.

Existing studies and present results indicate that cyclostomates have poorly developed musculature of the tentacle sheath, which is not surprising. In 1923, Borg reported no musculature in the tentacle sheath, although later he amended this statement, describing sparse and very delicate circular and longitudinal muscle fibers (Borg, 1926). Nielsen and Pedersen (1979) reported 15 strictly longitudinal muscles in *Crisia*. Unfortunately, Shunatova and Tamberg (2019) did not describe tentacle sheath in any of their studied species (*Tubulipora flabellaris, Patinella verrucaria and Crisiella producta*). In this study, examined species of hornerids also have very fine longitudinal musculature in the tentacle sheath.

The tentacle sheath and tentacles are the only portions of the cyclostomate polypide body wall where we find cellular and extracellular layers typical of the coelomates. In these regions the epidermis, ECM and coelothelium adhere to form a unified wall.

The digestive system of cyclostomates was examined by light microscopic methods primarily by Borg (1926) and later by Worsaae et al. (2018) and Schwaha (2018). At the EM level the primary source of ultrastructural information remains a study by Gordon (1975) on the gymnolaemate

### Cryptosula pallasiana

The general composition of the gut is typical for cyclostomates with a pharynx, cardia, caecum, pylorus and rectum. Unlike some bryozoans, e.g. *Zoobotryon verticillatum* (Bullivant & Bils, 1968), hornerids have no esophagus. Gut epithelium alternates between ciliated and non-ciliated, as well as bearing or lacking microvillar and cuticular cover. Cuticle is present in the mouth and pharynx only. Ciliation is present in the upper pharynx, and, more notably, in the pyloric region. The densely ciliated pylorus in particular resembles that of *C. pallasiana*. According to Gordon (1975), pyloric ciliature in this species generates a rotating rod which compacts the discarded caecum material. This cannot be checked in hornerids, however, because their opaque skeleton prevents observations of the digestive system at work.

Ciliation of the pharynx is somewhat irregular in hornerids: the number of cilia is unstable and all species except one have confirmed multiciliated myoepithelial cells. This is different from *Crisia eburnea* with a strict one-cell-wide vertical row of ciliated but non-muscular cells (Worsaae et al., 2018).

Typically for bryozoans, intracellular digestion occurs in the caecum; secondary lysosomes with undigested remains also accumulate in this region. Cells most vigorously engaged in biosynthesis are concentrated in the caecum and rectum, while myoepithelial cells of the pharynx mostly produce suction.

Overall, the pharynx of the studied hornerids is organized as a typical “self-contained” triradial suction pharynx (see Nielsen, 2013 for details) with myoepithelial walls. The same condition, reported previously for *Crisia* and *Cinctipora*, and now also for hornerids, is likely universal for Cyclostomatida.

The musculature of the hornerid digestive tract resembles that of other cyclostomates (Worsaae et al., 2018) and, to a degree, *C. pallasiana* (Gordon, 1975). The circular musculature of the mouth is continuous with that of the pharynx. The buccal dilators insert in the small gaps between the muscles of the oral sphincter and form a circle of radial spokes. The distinct circular musculature of the gut disappears past the cardia, whereas longitudinal muscles surround the entire digestive system. In contrast with the muscular mouth, no anal sphincter was seen.

The funiculus in cyclostomates is defined as a tubular peritoneal cord with longitudinal musculature (Schwaha et al., 2020). In *C. eburnea* (Worsaae et al., 2018), *C. elongata* (Carle and Ruppert, 1983), and the hornerid species examined here, there are two funicular muscles, whereas in *Cinctipora elegans* this number reaches eight (Schwaha et al., 2018).

Our results regarding the internal composition of the funiculus, however, differ from other existing accounts. Carle and Ruppert (1983) reported a fluid-filled lumen in the funiculus of *C. elongata*, and Schwaha et al. (2018) described it in *C. elegans* as filled with cells and lacking an opening (especially clear in Figure 5). Our findings are in marked contrast with both reports: in hornerids the funiculus contains no cells or openings, the entire core of the funiculus being a solid ECM cord.

The role of the funiculus in interzooidal connectivity is well-documented among gymnolaemates, but not in cyclostomates. The intriguing finding of mesenterial cells, connecting the funiculus laterally to the membranous sac and, through it, to the mural pores of *C. elongata* (Carle and Ruppert, 1983) remains unreproduced (see review by Schwaha et al., 2020). We also did not see any connections between the funiculus, the membranous sac and the interzooidal pores in any of the studied hornerids. For *Hornera robusta* in particular, a careful examination of the gap-free SBF SEM dataset confirmed that no such connection exists. The funicular attachment footprint measured in this study far exceeds the cross-section of the funiculus itself. It seems that high tensile stresses are transmitted through the funiculus and are distributed over a large area. We propose contractile function as primary for this organ in hornerids.

## Conclusions

The evidence from representatives of the previously unexplored suborder Cancellata strengthens and supports the unity of the general cyclostomate body plan, which is currently based mainly on examinations of the Crisiidae. At the same time, Horneridae have a few unique traits, including a lumen-free funiculus and an unbroken septum between lophophoral and trunk coeloms, which originates as lophophoral ECM and continues onto the outer surface of the pharynx. Some anatomical and morphometric differences between species and genera may support hornerid taxonomy, and a comparative analysis of polypide–skeleton interfaces at the attachment zones may provide insights into the biomechanics of cyclostomates, and, by implication, of some extinct stenolaemates.

## Acknowledgements

We are most grateful to Abby Smith (University of Otago) for providing organizational support and discussion, and to the master and crew of *R*.*V Polaris II*, staff at Portobello Marine Laboratory and the Otago Microscopy and Nanoscale Imaging unit (OMNI), University of Otago. Special thanks go to Hamish Bowman, Kim Currie, Linda Groenewegen, Allan Mitchell, Sharon Lequeux, Richard Easingwood, Amanda Fisher and Matthew Downes. Y.T. gratefully acknowledges support and discussion from Nina Alexeeva (Zoological Institute of the Russian Academy of Science) and funding from a University of Otago Doctoral Scholarship.

## Author Contributions

Y.T. designed the study, wrote the manuscript draft and prepared illustrations. Y.T. and P.B. sampled and processed bryozoans. R.N. provided financial and methodological assistance, and processed one of the SBF-SEM samples. Joint efforts went into discussion and editing of the manuscript.

## Data Availability Statement

The data that support the findings of this study are available from the corresponding author upon reasonable request.

